# Electrophysiological properties of the medial mammillary bodies across the sleep-wake cycle

**DOI:** 10.1101/2023.10.30.563083

**Authors:** Christopher M. Dillingham, Jonathan J. Wilson, Seralynne D. Vann

## Abstract

The medial mammillary bodies (MB) play an important role in the formation of spatial memories. Dense anatomical connectivity with hippocampal, brainstem, and thalamic structures, positions them as a focal point for the integration of movement-related and spatial information that is extended to the anterior thalamic nuclei and beyond to cortex. While their anatomical connectivity has been well-studied, much less is known about the physiological properties of the medial MBs, particularly in freely moving animals. We therefore carried out a comprehensive characterization of medial MB electrophysiology across arousal states by concurrently recording from the medial MB and the CA1 field of the hippocampus. In agreement with previous studies, we found medial MB neurons to have firing rates modulated by running speed and angular head velocity, as well as theta-entrained firing. We extended the characterization of MB neuron electrophysiology in three key ways: 1) we identified a subset of neurons (25%) that exhibit dominant bursting activity; 2) we show that ∼30% of theta-entrained neurons exhibit robust theta cycle skipping, a firing characteristic that implicates them in a network for prospective coding of position; 3) A considerable proportion of medial MB units show sharp wave-ripple (SWR) responsive firing (∼37%). The functional heterogeneity of MB electrophysiology reinforces their role as an integrative node for mnemonic processing and identifies potential roles for the MBs in memory consolidation through propagation of SWR-responsive activity to the anterior thalamus and prospective coding in the form of theta-cycle skipping.

**Significance Statement:** While the medial mammillary bodies (MBs) are important for memory, it is still not clear how they support memory formation. Through conjoint medial MB and hippocampal recordings across different arousal states we identified a population of medial MB units with diverse and often conjunctive physiological properties, including theta-entrained cells, cells modulated by running speed and angular head velocity, complex bursting, theta cycle skipping activity, and hippocampal sharp-wave ripple-responsive firing. These properties likely support a role for the medial MBs in mnemonic processing, enabling the integration of separate sensory streams and the propagation of information to the thalamus.

## Introduction

The medial mammillary bodies (MB) were one of the earliest regions associated with amnesia and play an important role in the formation of complex spatial memories, via their indirect influence on hippocampo-cortical networks (2009; Dillingham et al., 2015; Zakowski et al., 2017; Dillingham et al., 2019; McNaughton and Vann, 2022). While we have a detailed knowledge of the anatomical connectivity of the medial MBs, our knowledge of the electrophysiological profiles of the medial MBs in behaving animals is relatively limited. While a few studies have been carried out on *in vitro* preparations (Alonso and Llinas, 1992), or in anesthetized and/or head-fixed animals (e.g., Kocsis and Vertes, 1994; Bland et al., 1995; Kirk et al., 1996; Kocsis and Vertes, 1997) there is very little published data from awake, behaving rats (Sharp and Turner-Williams, 2005).

From these previous studies, we know that cells in the medial MBs show complex endogenous bursting activity (Alonso and Llinas, 1992; Kocsis and Vertes, 1994) and that a large proportion of neurons have spike discharges that are significantly phase-entrained to theta band oscillations (e.g., Bland et al., 1995). Additionally, firing rates of medial MB units have been reported to be highly correlated with both running speed and angular head velocity (Sharp and Turner-Williams, 2005). While, historically, the medial MBs have been considered simply a relay of hippocampal theta, the emerging picture appears more complex, with evidence suggesting a bi-directional influence between the hippocampus and medial MBs. Lesions (Sharp and Koester, 2008) or inactivation (Zakowski et al., 2017) of the mammillary bodies, or lesions of their primary efferent pathway, the mammillothalamic tract (Dillingham et al., 2019), alter the temporal dynamics of hippocampal processing, e.g., through attenuation of theta-band oscillatory frequency, highlighting the downstream impact of the MBs.

The present study sought to provide a comprehensive analysis of the electrophysiological properties of the medial MBs, extending beyond previous work by recording from greater numbers of neurons and, importantly, recording across different arousal states. A secondary goal was to assess the relationship between MB activity and hippocampal ripple-band activity. The medial MBs receive a dense input from the subiculum, via the postcommissural fornix, and the MB-projecting neurons in the subiculum are strongly modulated by hippocampal sharp-wave ripple events (Kitanishi et al., 2021). The prediction, therefore, is that MB activity will also be related to hippocampal sharp-wave ripple events. Given the strong links between the propagation of ripple activity, and memory, this may be an additional mechanism via which the extended hippocampal-mammillary network supports long-term memory formation.

## Methods

### Ethics

All experimental procedures were in accordance with the UK Animals (Scientific Procedures) Act, 1986 and associated guidelines, the EU directive 2010/63/EU, as well as the Cardiff University Biological Standards Committee.

### Animals

Eight male Lister-Hooded rats were used (Envigo, UK), weighing ∼300-350g at the time of surgery and behaviorally naïve. Prior to surgery, rats were group housed. Following surgery, implanted rats were housed singly (GR1800 double-decker cages; Techniplast, Kettering, UK) except for ∼1 hour daily, supervised socialization. Animals were maintained under diurnal light conditions (14 h light/10 h dark) with free access to water and environmental enrichment. When required, animals were food-deprived to no less than 85% of their free feeding weight to promote reward-based exploration.

### Electrode implantation

All surgeries were performed under an isoflurane-oxygen mixture (induction 5%, maintenance 2%–2.5% isoflurane) during the light phase of a 12 h, day/night cycle. Once anesthetized, animals were positioned in a stereotaxic frame (David Kopf Instruments). The skull was exposed and cleaned before 6-9 screws were secured and cemented to the skull. Craniotomies were made before careful removal of the dura and subsequent implantation of electrodes in positions corresponding to the following coordinates (mm from bregma unless stated): (CA1: AP: −3.8, ML: ±3.3, DV: −1.9 from top of cortex 4.4, ML: ±0.8, DV: −8.0 from top of cortex with a 4° angle toward the midline). Rats were implanted with 8 tetrodes (17 µm tungsten; California Fine Wire) into CA1, spanning approximately 1.4mm in the axial plane, and 4 octrodes (12 µm; California Fine Wire) into the mammillary bodies. Prior to implantation, electrodes were loaded into custom-made microdrives (Axona Ltd., St Albans) that allow for independent adjustment of electrode depth during recordings. Electrodes were then gold-plated (Gold plating solution; Neuralynx) to an impedance of ∼200-400kΩ using the open-source Tetroplater (https://github.com/MatsumotoJ/Tetroplater). Once lowered to the desired depth, a warmed paraffin/mineral oil (1:4) suspension, or synthetic dura (Dura-Gel; Cambridge Neurotech, Cambridge, UK) was applied to the dural surface and allowed to solidify or set, respectively. Drives were then secured in place with bone cement (Zimmer-Biomet; Swindon, UK). The scalp was then sutured around the drive. Post-surgery, animals were rehydrated with a subcutaneous 5–10ml injection of 4% glucose saline and given postoperative analgesia (0.06 ml, 5 mg/ml meloxicam, Boehringer Ingelheim).

### Behavior

Recordings were performed in a custom-made circular track arena (diameter 1.3 m) with a central sleep box (0.4 x 0.4 m). For awake (AWK) recordings, animals were trained to run outwardly around the track by accessing a small quantity of 50% water/condensed milk (by triggering one of two motion detectors positioned at 180° to one another) and returning to retrieve an experimenter-placed Cheerio (Nestle) reward at the start location. For sleep recordings, i.e., slow wave sleep (SWS) and rapid eye movement sleep (REM), animals were placed in a central sleep box with bedding and access to *ad libitum* food and water.

### Recordings

Recordings were performed during the light phase of a 12 h, day/night cycle following a postsurgery recovery period of no less than 8–10 days. Recordings were conducted using the DacqUSB acquisition system (Axona Ltd., St. Albans UK). Signals were amplified between 6000x and 12000x and bandpass filtered between 0.38 and 7 KHz for single unit detection. Spike detection and unit sorting was performed based on tetrode configuration of electrodes to remove duplicates within octrode recordings; units that shared >5% common spike times were identified and the cluster with the lower spike rate was discorded. Additionally, units with a firing rate of < 1 Hz were discarded. Local field potentials were sampled at a frequency of 4.8 kHz and downsampled to 2.4 KHz for all subsequent analyses. Automated spike sorting was performed using KlustaKwik (version 0.3.0: Kadir et al., 2014) and manual verification and adjustment was performed using Tint (Axona Ltd., St Albans, UK).

### Perfusion/histology

At the end of experiments, animals were given an overdose of sodium pentobarbital (60 mg/kg, Euthatal, Rhone Merieux) and transcardially perfused with 0.1 m PBS followed by 2.5% PFA in 0.1 m PBS. Brains were removed and postfixed overnight in 2.5% PFA before being transferred to 20% sucrose in 0.1M PBS for approximately 24 hours. Sections were cut at 40 μm in the coronal plane using a cryostat. A 1-in-3 series was collected directly onto gelatin-coated slides and Nissl-stained (cresyl violet, Sigma-Aldrich) for verification of electrode location.

### Data analyses

All analyses were performed using custom scripts in MATLAB (version 2022a; The MathWorks) and Python 3.6.6, unless otherwise stated.

### LFP analyses

POWER SPECTRAL DENSITY/COHERENCE: A multi-taper method (Bokil et al., 2010) was used for calculation of frequency domain spectral power, and the coherence between signals was estimated using magnitude-squared coherence. Power spectra were normalized to account for differences in electrode impedance by the integral of the power in the spectrum. Optimal LFP channels for use in analyses across a given session was chosen based on a combination of the spectral power in the theta band (4-12 Hz) during AWK recordings (MB and HPC), as well as the number of SWRs recorded during sleep recordings in the same recording day (HPC only).

GRANGER CAUSALITY: In order to estimate the strength and directionality of information transfer between the medial MB, and the hippocampal LFP, we employed the MVGC toolbox, which utilises the Weiner-Granger causality method (Barnett and Seth, 2014). This method assesses the degree to which the information content in one time domain signal predicts another in the frequency domain using autoregessive models. Local field potentials from both brain regoins were input as unfiltered signals, downsampled to a sampling frequency of 120 Hz. The model order was determined using the Levinson-Wiggins-Robinson algorithm and a maximum model order was defined as 50 sample points and the optimal order was defined using the Baysien Information Criterion (BIC). Peak Granger Causality values that were significant at an alpha of p< 0.001, as defined by Granger’s F-Test were compared in the theta band (6-12 Hz) across subjects.

SLEEP DETECTION (SWS/REM): In the hippocampus, the awake state is characterized by high theta, and low delta power. To detect episodes of SWS, the short time Fourier transformation of the signal was first calculated using the multi-taper method (http://chronux.org; Mitra and Bokil, 2007) with a 10 s moving window and 4 s overlap. A theta/delta ratio for each time window was calculated using a peak detection algorithm on a smoothed (moving-average) of the maximum-normalized and squared signal of the spectrum. A global (whole recording) manual threshold based on visual classification was used to achieve optimal separation of AWK and NREM states (a lower than threshold TD ratio was classified as NREM and higher than threshold, AWK). Similarly, REM states were identified using the theta/delta ratio, with a speed threshold of 4 cm/s set to separate REM and AWK states.

SHARP WAVE RIPPLE DETECTION: Candidate SWRs were detected using peak detection threshold of the mean plus four standard deviations of the 150-250Hz bandpass-filtered SWS-indexed hippocampal LFP signal, with a minimum inter-peak time period of 200 ms. From ripple triggered averages of candidate SWR times (20 ms), the number of oscillations above the standard deviation threshold were counted and SWR events were considered valid if there were at least 5 oscillations above threshold within a 20 ms window surrounding the putative event.

REM-SWS COMPARISONS: To assess changes in firing activity during different sleep states, REM and SWS epochs were first defined. REM events, identified as timepoints with high theta/delta power ratios and sub-threshold (<4 cm/s) movement speeds, were concatenated to define single REM epochs if there were fewer than 60 s between REM events. SWS epochs were defined similarly; SWS intervals were concatenated to form SWS epochs if there were fewer than 60 s between SWS events. For each day where there were identifiable REM epochs of over 30 s duration, the REM epoch with the longest duration and with neighboring SWS epochs was selected for analysis. Mean firing rates were calculated for MB and putative HPC pyramidal cells during REM and the neighboring SWS epochs. Firing rate scores for each sleep state were calculated as:

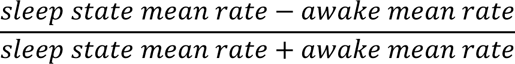

Units that did not fire in any of the sleep states were excluded from firing rate analysis.

Cell-pair correlations were calculated for MB units and HPC pyramidal units separately. Correlations were performed on single unit spike trains binned into non-overlapping 100ms bins. Cells that were inactive in any of the three sleep states were excluded from analysis. Synchrony was defined as the mean within-region cell-pair correlation for each sleep state epoch. Two-way repeated measures ANOVA were used to compare firing rates and synchrony across regions and sleep states. Post-hoc pairwise t-tests were Bonferroni corrected for multiple comparisons.

### Single unit analyses

AUTOCORRELATION: Autocorrelation histograms were calculated as the cumulative sum of spikes found within ±0.5 seconds of each spike. Spike train autocorrelations were normalized so that the cumulative sum of 1 ms bins equal 1.

RUNNING SPEED MODULATION: Running speed was derived from two-dimensional positional data (sampled at 10 Hz), using a ceiling-mounted infra-red video camera, which in turn detected a small and a larger LED cluster mounted on the headstage. Positional information was resampled to 2.4 KHz and smoothed using a moving average filter across 100 ms windows. Spike rates were calculated for 1 cm/s bins (range 1-60 cm/s). Speed bins were included only if the range had been sampled for a minimum of 1 s within the recording session. Spike counts per cycle were calculated by first detecting the peaks of 4-12 Hz bandpass filtered hippocampal LFP. The integer rounded mean running speed of each theta cycle and the number of spikes for each corresponding cycle were calculated, allowing the mean average number of spikes per cycle in each speed bin (1-60 cm/s) to be determined. To assess the relationship between phase preference and running speed, speed values for each spike were calculated by finding the spike triggered average of the running speed (200 ms windows). Spike phase values were calculated through piecewise cubic polynominal interpolation of phase angle values of the hippocampal LFP. Mean resultant vector lengths (MRV; square root of the sum of the sin and cosine components of the phase values) of phase values from each valid speed bin were calculated. For each unit, the relationship between firing rate and running speed was assessed using linear regression. Where significant relationships between running speed and firing rate were observed, likelihood ratio tests were implemented to determine whether the inclusion of a quadratic term in nested models significantly improved the model fit (p<0.05).

WAVEFORM: Waveforms were extracted for each unit (50 bins of 0.04ms) from the recording channel with the largest mean peak waveform amplitude. Waveforms were normalized by peak firing amplitude and any inverted waveforms were excluded from analysis. Putative hippocampal interneurons and pyramidal neurons were identified through KMeans clustering of the first two principal components of hippocampal unit mean normalized waveforms and the z-score of their mean firing rate (English et al., 2017; Nitzan et al., 2020). Putative interneurons were excluded from subsequent analysis. Waveform asymmetry was defined as the difference between the minimum amplitudes before and after the peak. Waveform peak-trough width was defined as the number of bins between the peak and the minimum amplitude after the peak. Peak-trough width and asymmetry measures were compared between MBs and HPC using MANOVA and post-hoc one-way ANOVAs.

ANGULAR HEAD VELOCITY: Angular head velocity (AHV) was calculated as the first derivative of head direction based on the first five minutes of each central sleep box recording. AHV calculations were based on recordings of quiet wakefulness in the sleep box, rather than the circular track, to remove the potential confound of running speed-dependent changes in firing rate derived from changes in heading direction that naturally occur when running in a circle. First, head direction (HD) was derived from the X and Y coordinates of one small and one large LED cluster that were mounted on the headstage. Head direction data were resampled to 1KHz and smoothed using a moving average filter across 100 ms windows. To obtain corresponding AHV values, a moving window (400 ms) surrounding each sample was indexed and the linear slopes (AHV values) were calculated. AHV values were assigned to 0.1 radians/s bins within a range of ±5 radians/s; bins with less than 1 s of occupancy were discarded. Firing rates within each bin were then calculated and tuning curves were generated for each unit. For each tuning curve, linear regressions were performed, separately, on clockwise (-5 to 0 radians/s), and counterclockwise (0-5 radians/s) head movements. A shuffling procedure was then performed, in which spike trains were circularly shifted (500 iterations) by randomly generated values between 20-100 s. Thresholds were then calculated from distributions of the estimates derived from linear regression of the shuffled-generated tuning curves. A unit was considered to be significantly AHV modulated if the estimate generated from non-shuffled data was greater than the upper bound (99% confidence interval) of the shuffle distribution (clockwise movement), or lower than the 1% confidence interval of the shuffle distribution for counterclockwise head movements.

THETA RHYTHMICITY/PHASE: The theta index of a unit, as defined by the ratio of the power of the autocorrelogram within 1 Hz of the theta peak and the power within the entire spectrum of the autocorrelogram (Climer et al., 2015), provides a measure of the degree to which a unit engages in theta rhythmic firing. To assess theta cycle phase preference, a Hilbert transformation was applied to the 4-12 Hz bandpassed filtered (3^rd^ order Butterworth) hippocampal LFP. Spike phase values were calculated through piecewise cubic polynomial interpolation of the LFP instantaneous phase values. To classify units as phase entrained, phase tuning curves were generated from circularly shifted spike trains (1000 iterations, shifted by a random value between –5 and 5 s). The mean resultant vector (MRV) length was calculated for the shuffled and non-shuffled data and a unit was classified as significantly phase entrained if the non-shuffled MRV was greater than the 95^th^ percentile of the distribution of MRVs based on shuffled spike trains.

BURSTINESS: The burst probability for each unit was calculated as the number of bursts divided by the sum of the number of bursts and spikes in the session. Burst detection was performed according to the method of Oosterhof and Oosterhof (2013). Bursts were defined as complexes with a minimum of 3 spikes, with a maximum ISI to start a putative burst of 6 ms, and a maximum continuing ISI of 8 ms. Medial MB units were classified as dominant bursting (DB) or sparsely bursting (SB) based on the methods of Simonnet and Brecht (2019). Specifically, principal component analyses were performed on the probability normalized log ISI histograms (logISI; 0.001-10 s), and the first 20 ms from the central peak of the autocorrelograms for each unit. Ward’s hierarchical clustering method was subsequently performed on the first principal component of the autocorrelation and the first to third principal components of the logISIs. Principal component inclusion was based on the variance explained and the visual inspection of population logISIs and autocorrelations when sorted by PCA score.

SWR-RESPONSIVE ACTIVITY: To assess the degree to which medial MB unit firing was positively-, negatively-, or unmodulated by hippocampal SWRs, spikes within ±500 ms of each SWR were summed to generate peri-SWR time histograms (PSTH) for each MB, or HPC unit. To determine the significance of responsive firing, PSTHs were generated from shuffled datasets in which ripple event times were circularly shifted (1000 iterations) by a random set value of between –5, and +5 s. An SWR responsiveness index (SWR RI) was calculated for each true PSTH by subtracting the mean firing rate from a 250 ms window around the shifted event time from the same period of each actual PSTHs. From each shuffled PSTH, the mean firing rate within the same window of the remaining 999 shuffled PSTHs were subtracted to generate a normal distribution of SWR responsiveness indices. The values of the SWR responsiveness indices of the non-shuffled PSTHs were compared with respect to the 5th and 95^th^ percentiles of the shuffled distribution in order to assess the likelihood that the observed peri-SWR firing could happen by chance.

### Statistical analyses

Statistical analyses were performed either in R Statistics (version 4.3.1; provided in the public domain by the R Foundation for Statistical Computing, Vienna, Austria; R Development Core Team, 2009, available at http://www.r-project.org/), in MATLAB (version 2022a; The MathWorks), or in python (version 3.6.6 – using the packages *pingouin* and *statsmodels*). For within unit analyses of firing rate correlates of movement, i.e., running speed and angular head velocity, linear models were fitted. For analyses in which parameters were compared between assigned groups, e.g., DB vs SB, a random term (1|subject ID) was included in linear mixed effects models to account for the assignment of units to different animal subjects. Given the difficulties associated with determining degrees of freedom in these instances, p values were obtained through likelihood ratio tests comparing nested models. Unless otherwise stated, the threshold for significance was set at p<0.01. Graphs were generated using either Matlab, the python package *seaborn*, or the “ggplot2” package in R Statistics (Wickham, 2016), while figures were compiled in Inkscape (Inkscape version 0.92.4, The Inkscape Project, freely available at www.inkscape.org).

## Results

Eight rats were implanted with octrodes and tetrodes targeting the medial MBs and the CA1 field of the hippocampus (HPC), respectively. Following histological assessment, 6 cases were included for subsequent analyses (**Fig. 1A**). In two cases (MamHPC10, and 11, the paths of the electrodes were located medially within pars medialis of the medial MBs. In the remaining four cases, the electrode paths (range: 700-1400 µm paths; **Fig 1A**) were along or close to the pars medialis/pars lateralis boundary. Across all recording sessions (range 10-14 recording days/rat), 543 single units were recorded from octrodes in the medial MBs while rats were running on a circular track. Of these, 293 units were recorded from whilst rats were in the central arena, which included quiet wakefulness and slow wave sleep (SWS), and rapid eye movement sleep (REM; n = 145 units). Across the same recording sessions, 211 units were recorded from tetrodes implanted in the HPC (PYR: n = 179; INT n = 32) during wakefulness, and 110 HPC PYR units were recorded during REM/SWS.

**Figure 1.**
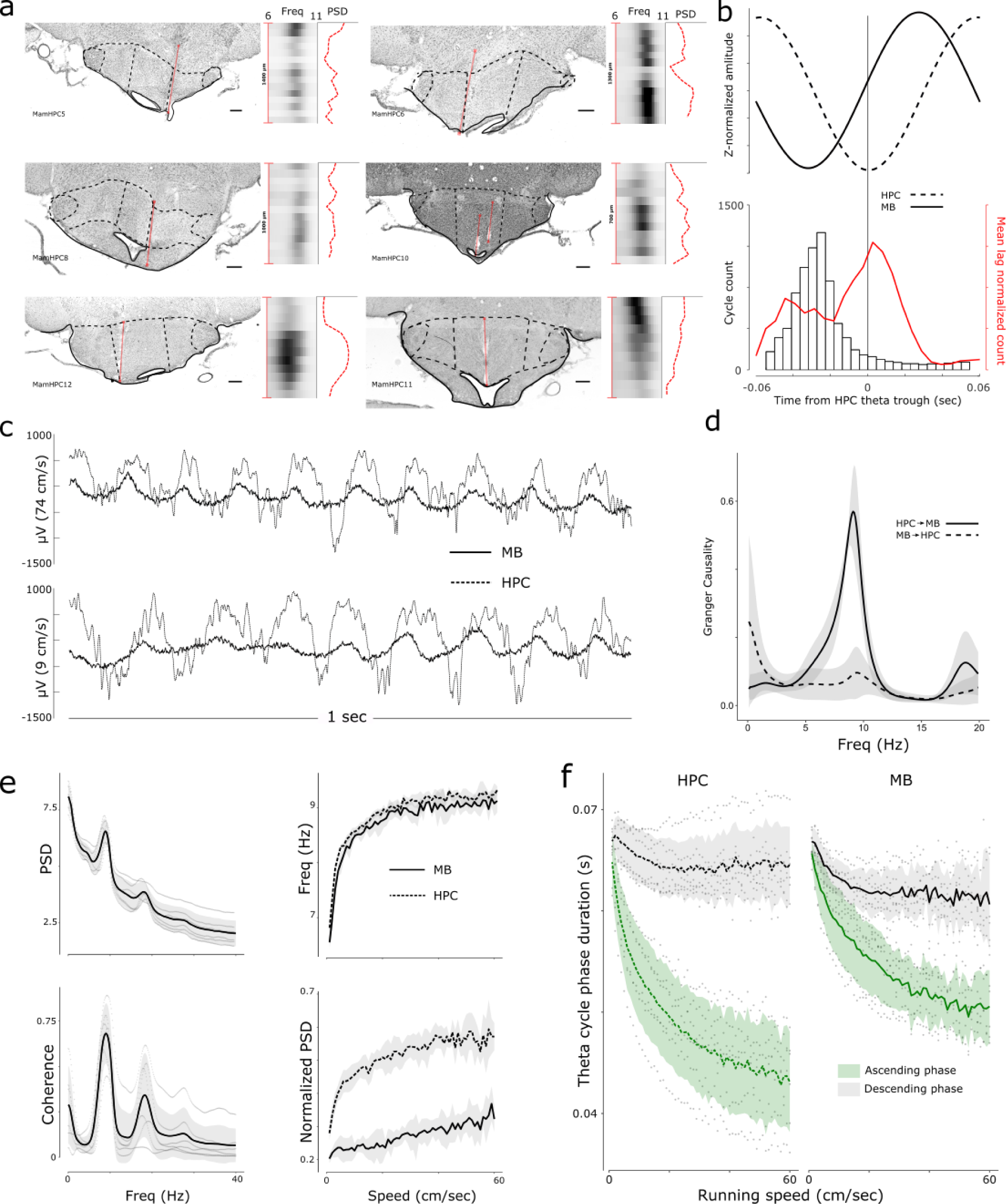
Local field potential (LFP) characteristics of the mammillary bodies (MB) from conjoint hippocampal (HPC), and medial MB recordings. **A**, Histological reconstructions of the MB electrode tracks (red lines). Black and white heatmaps show the theta band (7-10 Hz) power of the LFP through the depth of the electrode track. Associated red dotted line shows the peak LFP power of the LFP vs. electrode depth; **B**, Mean phase lag in an example awake session between HPC (dashed) and medial MB (solid) LFPs showing ∼30 ms lag (upper), and a histogram of lags cycle lags within the session (lower). Red trace shows the mean lag between the LFPs from all animals combined; **C**, Example LFP traces, from conjoint MB and HPC recordings. Top traces: mean running speed of 74 cm/s; bottom traces: 9 cm/s illustrating change in frequency and cycle asymmetry. **D**, Granger causality for HPC-to-MB and MB-to-HPC during active wakefulness; **E**, Power spectral density (PSD) of the MB LFP showing peaks in the delta, theta, and first harmonic of theta **F**, Comparison of ascending (trough-to-peak) and descending (peak-to-trough) theta cycle durations with running speed. All scale bars in **A** are 500 µm.

### Local Field Potential characteristics

Local field potentials (LFP) are thought to represent the post-synaptic changes in membrane potential around the recording electrode. Given that hippocampal (subicular) efferent projections represent a major input to the MBs, we first wanted to establish the extent to which the characteristics of the LFP recorded from the medial MB mirrored those recorded from CA1.

The power spectra of LFP of the electrodes placed in both CA1 and in the medial MBs had a dominant peak in the theta band (6-12 Hz), accompanying delta (1-4 Hz) peaks, as well as peaks in the beta (12-20 Hz) band, corresponding to the 1^st^ harmonic of theta. During awake periods (AWK), LFP signals from the two brain regions showed high peak magnitude-squared coherence in the theta frequency band (6-12 Hz; 0.693±0.154). As well as being highly correlated in terms of change in amplitude, one would predict that the unidirectional nature of the HPC-MB projection (Meibach and Siegel, 1977) would be reflected in a phase lag in theta oscillations between the HPC and medial MB signals. While the mean lag between the signals was consistent with the anatomy 0.012±0.006 sec; **Fig. 1B**), there was considerably more variance both between subjects, and within recording sessions than might have been predicted. Theta frequency in the hippocampal and extra-hippocampal limbic system increases as a function of running speed (Sławińska and Kasicki, 1998; Maurer et al., 2005; Jeewajee et al., 2008; Alex et al., 2016; Carpenter et al., 2017), thus we next looked at whether some of this variance in phase lag could be explained by differences in their relationship between phase lag and running speed. No relationship between phase lag and running speed was apparent (-1.17e^-6^ ± 1.42e^-5^, t=-0.08, Chi = 0.007, p = 0.94), and consistent with this finding, peak theta frequency in the LFPs of both regions showed a significant theta-cycle symmetry-dependent association with running speed (MB quadratic fit: -0.001 ± 5.114e^-5^, t=-18.08, p<0.001; HPC quadratic fit 8.364e^-4^ ± 4.186e^-5^, t = -19.98, p<0.001; **Fig. 1E-F**) that did not statistically differ from one another (0.001 ± 0.001, t=-0.665, p=0.506). While phase lag between signals did not present a clear picture of the direction of flow of information between HPC and medial MB, Granger causality analysis of the synchronously recorded signals was more conclusive, with the magnitude of the theta band Granger causality score in the HPC-MB direction significantly higher than that in the inverse direction (0.476 ± 0.058, t = 8.25, p<0.001; **Fig. 1D**).

### Medial mammillary body unit characteristics

The characteristics of medial MB neurons of rats in an awake state were found to be highly consistent with those reported in the work of Sharp and Turner-Williams (2005). The majority of units recorded fired either as a function of running speed and/or angular head velocity and/or fired at a theta-band frequency with phase entrainment to hippocampal theta oscillations. In addition, and consistent with studies in both slice (Llinas and Alonso, 1992) and anesthetized preparations (Kocsis and Vertes, 1994), we also found approximately a quarter of recorded medial MB units to exhibit complex bursting activity.

### Medial mammillary body unit waveforms

Neurons in the medial MBs constitute a population of excitatory projection neurons, that predominantly project via collaterals to the anterior thalamic nuclei and Gudden’s ventral tegmental nuclei (VTg; Van Der Kooy et al., 1978; Ching Liang, 1983; Vann et al., 2007). We compared the waveform shapes of medial MB units with those of HPC pyramidal (PYR) units, which have characteristically broad and asymmetric waveforms (Csicsvari et al., 1998b). Both peak-trough width and waveform asymmetry differed between MB units and putative excitatory PYR units (Pillai’s trace = 0.73, *F*_(2,95)_ = 126.79, p <0.001), with post-hoc comparisons revealing MB units are narrower (MB: 0.4 ± 0.01ms; PYR: 0.74 ± 0.03ms; *F*_(1,96)_ = 154.85, p<.001) and more symmetrical (MB: -0.13 ± 0.02; PYR: -0.44 ± 0.03; *F*_(1,96)_ = 95.6, p <.001) than PYR units (**Fig. 2D**).

**Figure 2.**
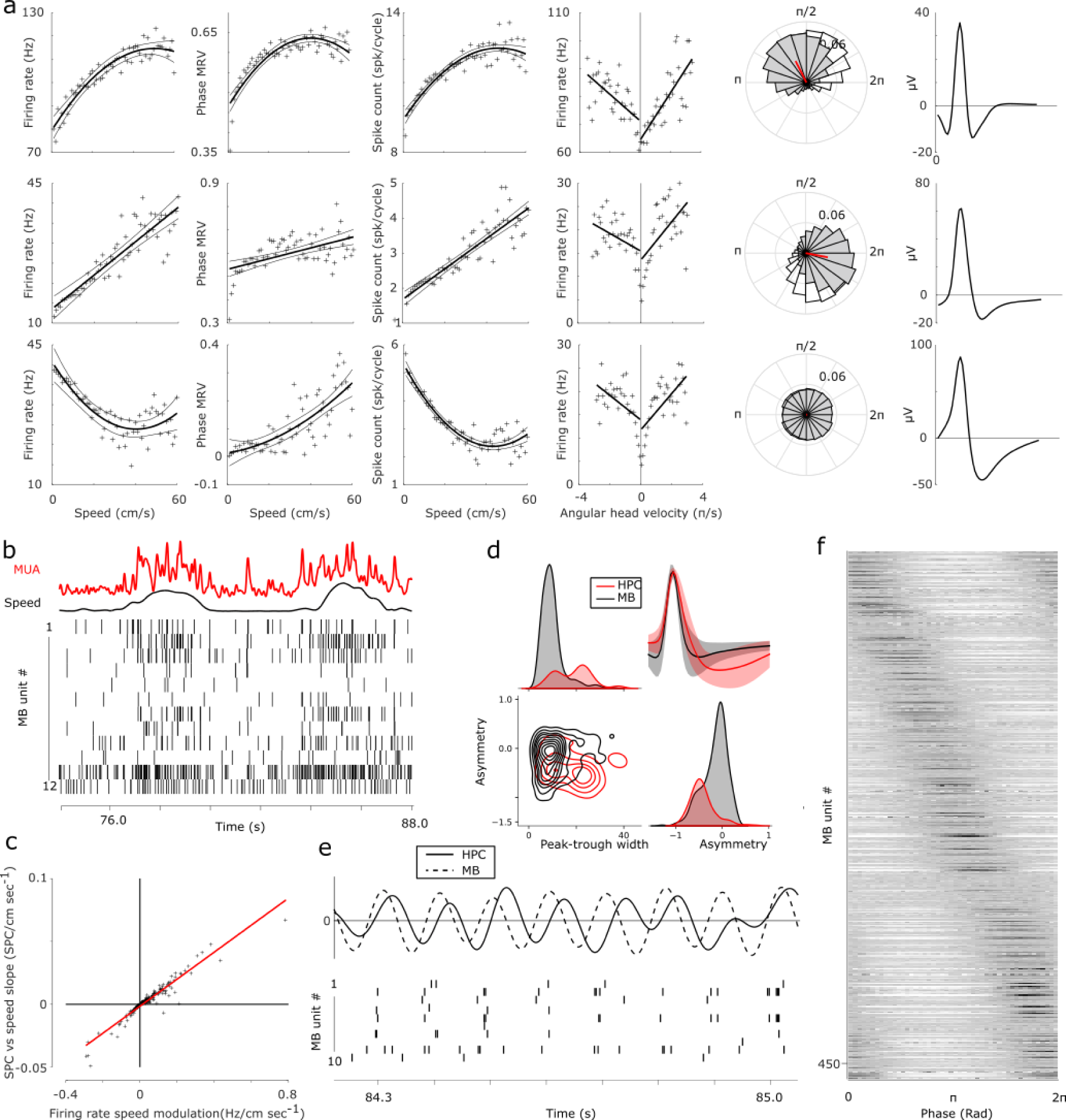
Physiological characteristics of mammillary body units during awake exploration. Each row in **A** depicts the speed-, angular head velocity- and theta phase-dependent firing properties, as well as spike waveforms of representative medial MB units. Alongside speed/firing rate correlations, medial MB units also exhibited significant changes in strength of phase entrainment with running speed; **B**, Example raster plots of synchronously recorded medial MB units showing population-dependent firing with respect to running speed. Red trace shows the combined multi-unit activity (MUA) of the units depicted in the raster plots. **C**, As demonstrated in the examples in **A**, a unit’s firing rate by running speed-, and its spikes per cycle by running speed relationship were highly correlated across the population of medial MB units recorded, reflecting an active increase in firing rate, i.e., an increase in spikes/cycle; **D**, Waveform asymmetry and peak-trough width of medial MB units (black) and hippocampal PYR units (red). Mean spike waveforms of medial MB (black) and hippocampal (red) units (top-right); **E**, example epoch of synchronously recorded medial MB (dashed), and HPC (solid) local field potentials with corresponding raster plots of medial MB units exhibiting phase entrainment of firing during awake exploration; **F**, Population phase preference of medial MB units (sorted by preferred phase), with each row representing the spike count phase histogram of a single medial MB unit.

### Running speed-correlated firing

The firing rates of a large proportion of units (341/543 units; 62.8%) were found to be significantly correlated with running speed (**Fig. 2A-B**). By evaluating whether the inclusion of a quadratic term significantly improved the fit of models, units were classified as either linearly (n = 227/543; 41.8%), or non-linearly correlated (n = 114/543; 21.0%). Of those 227 units that were linearly correlated with running speed, 152 (67.0%) were positively correlated (**Fig. 2A**; upper and middle example units), while of the 114 units that were best fit by a quadratic term, 68 units (59.7%; e.g., **Fig2A**, lower example unit) were positively correlated. As has been described in the medial entorhinal cortex (Hinman et al., 2016), a proportion of conjunctive speed correlated firing and spike phase entrained medial MB units (110/543; 20.3%), were found to have significant correlations between speed and the strength of phase entrainment, i.e., the mean resultant vector length of medial MB spike phase histograms was correlated with running speed. In order to exclude the possibility that firing rate/running speed relationships were not simply a function of phase-entrained units firing at a higher rate due to an increased number of theta cycles per second (Hinman et al., 2016), we examined the relationship between running speed and spikes per theta cycle in medial MB units. Consistent with an active increase in firing rate with running speed, regression analysis showed a highly significant positive relationship between running speed/firing rate and running speed/spikes per cycle (estimate value: 0.106±0.002, t = 53.65, adjusted R^2^ = 0.886, p<0.001; **Fig. 2C**).

### Angular head velocity correlated firing

Angular head velocity (AHV) tuning has been reported previously in the medial MBs (Sharp and Turner Williams, 2005). To determine whether the firing rates of medial MB units varied as a function of AHV, we analysed five-minute periods of quiet wakefulness in the central sleep box rather than the circular track which could have artificially affected lateral head movements. Linear regression performed on clockwise, and counter-clockwise head movements independently, revealed that 23.2% (68/293) of medial MB units showed significant angular head velocity tuning in both directions, while an additional 30.0% (88/293) units showed asymmetrical tuning in either the clockwise (60/88 units), or counter-clockwise (28/88 units) direction.

### Theta phase entrainment

Perhaps the most well-defined physiological marker for recordings in the medial MBs is the abundance of theta entrained units. Nearly 50% (267/543) of medial MB units were found to fire rhythmically within the theta frequency band. In turn, we found that more than 85% (463/543) of units showed significant phase preference within theta (6-10 Hz) cycles (**Fig 2A, E**). Phase preference of medial MB units was distributed uniformly across the theta cycle both as a whole population, or when indexed based on conjunctive firing properties, e.g., phase entrained, speed units (n = 543 medial MB units: Rayleigh Z = 0.47, p=0.63; **Fig. 2A; 2F**). Approximately 55% (299/543) of medial MB units were both phase entrained and responsive to running speed while nearly 30% (159/543) of medial MB units showed both theta rhythmicity in their firing, and speed-dependent activity.

### Bursting

An additional characteristic of AWK medial MB unit activity was the observation of bursting activity. We employed the approach of Simmonet and Brecht (2019) to aid the classification of bursting units. For this, we performed hierarchical clustering on PCA scores, calculated from the log inter-spike interval (logISI), as well as the 0.02 second lag of the autocorrelogram of all medial MB unit spike trains (**Fig. 3A**). Employing the same terminology as that used by Simmonet and Brecht in the subiculum, clustering of MB units identified two distinct groups: sparsely bursting units (SB; 409/543 units; 75.3%), and dominantly bursting units (DB; 134/543 units; 24.7%; Fig. 3A). To determine the effectiveness of this classification, a comparison of unit burst probability between the clusters revealed that DB units had a significantly higher propensity to burst (difference in burst probability: 0.07±0.002, t = 24.10, Chi = 408.84, p<0.001; **Fig. 3C**), while firing rates were also significantly higher in the DB group (difference between DB and SB: 20.94±1.12 Hz, t = 18.62, Chi = 268.92, p<0.001; **Fig. 3D**). Furthermore, visual inspection of the logISIs of DB and SB clusters (**Fig. 3B**; black and red, respectively) showed a LogISI peak in the DB cluster, spanning ∼2-15ms that was largely absent in the SB cluster, reflecting the short latency between spikes within a burst event. Similarly, high spike counts within a ∼5ms lag of the DB autocorrelograms (**Fig. 3C**; black), that were not apparent in the majority of units belonging to the SB cluster (**Fig 3C**; red), is reflective of a relative absence of spike activity within 0-10 ms of preceding activity in SB units.

**Figure 3.**
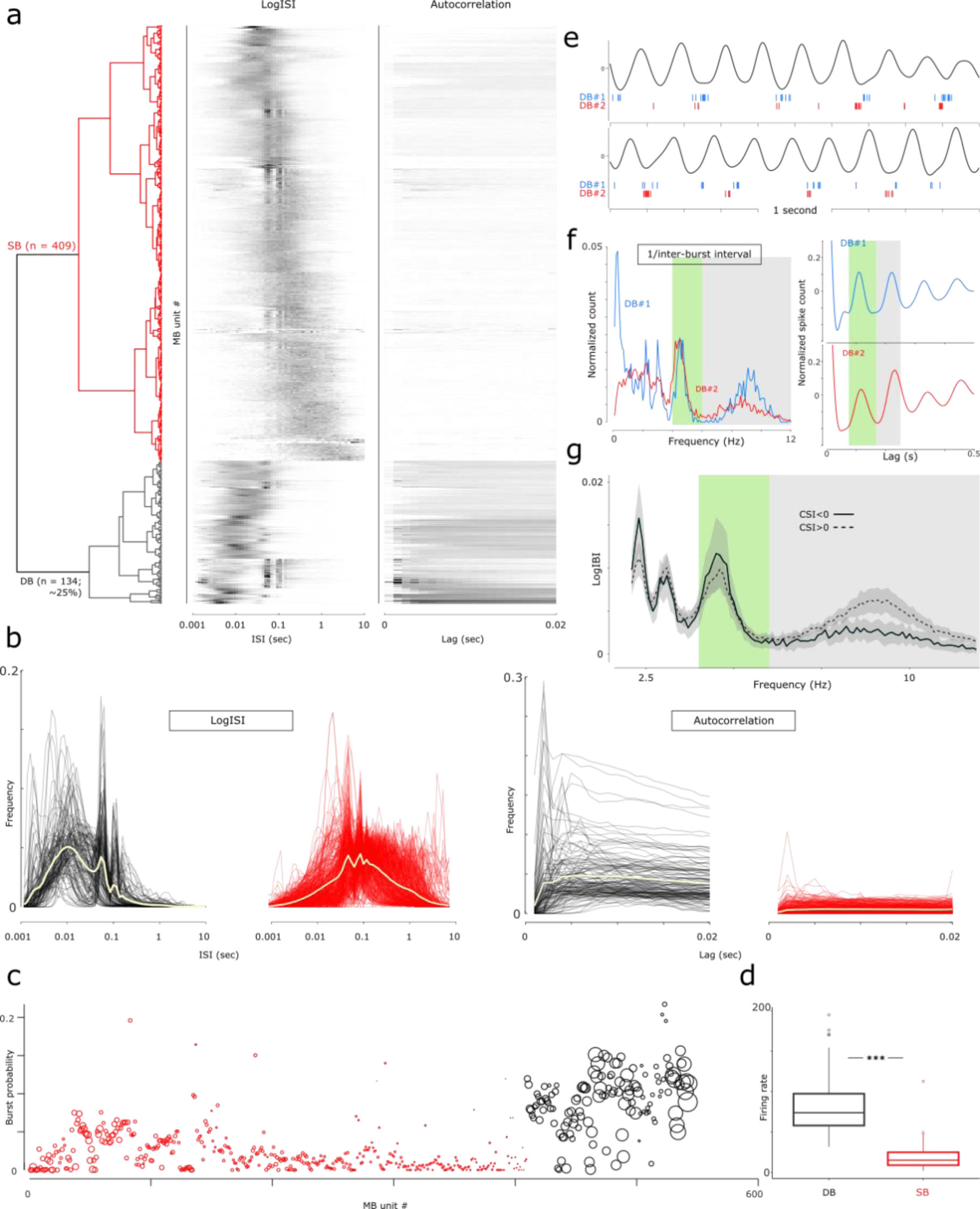
Bursting neurons in the medial mammillary bodies (MBs). Using the approach of Simonnet and Brecht (2019), PCAs 1-3 of the logISI and the first PCA of the 0.02 second autocorrelation explained 80.1% and 87.0% of the variance, respectively. **A**, Ward’s agglomerative clustering identified two clusters (dendrogram) equivalent to the dominant bursting (DB; n=134; 24.7%; black) and sparsely bursting (SB; n = 409; 75.3%; red) of Simonnet and Brecht (2019); Neighboring heatmaps show log inter-spike interval histograms (LogISI; middle), and autocorrelations (maximum 0.02 s lag), ordered according to the clustering in **A**. In each case a row corresponds to a single medial MB unit. **B**, left: Medial MB, DB units (black), had a higher mean peak in the logISI around 4-15 ms and a secondary peak shared by SB units (red) at ∼100 ms, corresponding to firing at a frequency within the theta band; right: the mean 0.02 s autocorrelogram of DB units (red) showing a higher initial peak than SB units (black). The overlaid yellow trace in the four panels of **B** show the mean averages for each case; **C**, A scatter plot showing the burst probability of medial MB units, sorted according to the hierarchical clustering procedure, is higher in DB classified units (black) in **A**. Circle size represents unit mean firing rate. **D**, Boxplot showing that as a population, firing rates in DB units are significantly higher than SB units. **E**, Two example bursting medial MB units (DB#1, blue; DB#2; red) exhibiting cycle-skipping bursting (DB#1 in upper example and DB#2 in the lower trace), and non-skipping bursting activity (DB#2 in upper trace, and DB#1 in lower example), at different points within the same recording. **F,** Left: the corresponding log 1/inter-burst interval histograms and theta-bandpass filtered 0.5 second autocorrelograms of example units DB#1 and 2. Peaks between 4-6 Hz correspond to cycle-skipping bursting activity, while the peaks at 6-12 Hz correspond to cycle-by-cycle bursting. Right: autocorrelograms for both example units show theta rhythmicity, while that of DB#2 shows a characteristic theta-skipping trace, likely to correspond to more dominant cycle-skipping burst activity; **G**, Line plot showing the mean average log inter-burst interval histogram of all medial MB units. Prominent peaks within the theta (6-12 Hz; green) and low theta (gray; 4-6 Hz) are highlighted.

Using this classification for DB units, we next assessed the degree to which bursting activity was associated with theta-dependent-, running speed- and angular head velocity-dependent-unit firing. Subgroups of units exhibiting conjoint bursting/speed represented 83.6% of (112/134) DB units; DB/theta rhythmicity in spike trains represented 50.8% (68/134) of DB units, and 82.1% (55/67) DB units also fired as a function of angular head velocity. Finally, units that exhibited combined bursting, speed-dependent firing, and theta rhythmicity in firing accounted for 46.2% of the DB units.

### Theta cycle skipping

Closer inspection of longer-duration (0.5 s lag) autocorrelograms of medial MB units revealed characteristic examples of theta-skipping rhythmicity, i.e., in which the count at the secondary peaks at around 0.2-0.3 s lag was greater than the primary peak at 0.1-0.15 s (e.g., Climer et al., 2015). We next classified putative theta-skipping units by calculating the cycle skipping index (CSI; Kay et al., 2020), which defines the relative proportions of primary and secondary peaks of the autocorrelograms of theta rhythmic activity for each MB unit with a significant theta index. Of 267 theta rhythmic medial MB units, 80 (30%) had a CSI greater than zero. Of these 80 putative theta-skipping medial MB units, 20 (25%) were also classified as DB units, which was representative of the proportion of DB units within the total medial MB population. To investigate the occurrence of theta cycle skipping DB unit firing, we calculated log inter-burst interval (LogIBI) histograms for all DB-classified medial MB units. Across all DB units, the mean LogIBI histogram had two dominant peaks, the first within typical theta range (6-12 Hz) and a second at approximately half this frequency (4-6 Hz). Consistent with theta-skipping DB-activity in these 20 medial MB units, relative proportions of theta and half theta bursting LogIBI peaks were highly correlated with the CSI (6.15±0.53, t = 11.55, p <0.001). Spike rasters relative to the theta bandpassed HPC LFP showed that single spike, or burst activity, cycled between epochs in which spike/burst events were observed in each theta cycle, and those in which such events were observed only every other cycle (**Fig. 3F-F^ii^**).

A comparison of the LogIBIs of DB units with CSI values < 0 (i.e., theta DB units) and those with CSI values >0 (i.e. putative theta skipping DB units) showed that the integrated amplitude of the LogIBI in the theta band (6-12 Hz) of non-skipping (CSI<1) DB units was significantly greater than that of theta skipping DB units (non-theta skipping vs. theta skipping units: 0.14±0.05, t = 2.65, Chi = 6.71, p = 0.010; **Fig. 3G**). Interestingly, however, both classifications had a comparable bursting count peak in the 4-6 Hz (half-theta) band, suggesting that while the majority of DB units exhibited theta skipping DB activity, the relative proportion of theta versus theta-skipping bursts is variable within the population (e.g., **Fig. 3F^i^**).

### Physiological changes in the medial mammillary bodies during sleep

The rodent sleep cycle can be broadly split into awake (AWK), slow wave sleep (SWS), and rapid eye movement (REM) phases. Unlike AWK and REM phases, which are dominated by theta band activity, SWS is characterized by delta band (1-4 Hz) slow waves, which are highly synchronous across hippocampo-cortical regions. Similarly, the medial MB LFP is also characterized by delta band oscillations during SWS (medial MB: 1.25 ± 0.13 Hz; HPC: 1.57 ± 0.07 Hz).

As found during AWK periods, the multi-taper power spectra of the medial MB LFP during REM showed dominant peaks of power in the theta band (medial MB: 7.80 ± 0.28 Hz; HPC: 7.76 ± 0.25 Hz; **Fig. 4A**). Consistent with the known HPC-medial MB anatomical connectivity, the magnitude-squared coherence of HPC and medial MB signals was high in the theta frequency band during REM (coherence: 0.557 ± 0.056; peak coherence frequency: 8.203 ± 0.397 Hz; **Fig. 4A**). Again, similar to the findings in AWK (**Fig. 4A**; also see **Fig. 1D**), the magnitude of theta band Granger causality scores was significantly higher in the HPC-MB direction than the inverse in REM (Granger score HPC-to-medial MB vs medial MB-to-HPC: 0.887 ± 0.170, t = 5.216, p = 0.002; **Fig. 4A**).

**Figure 4.**
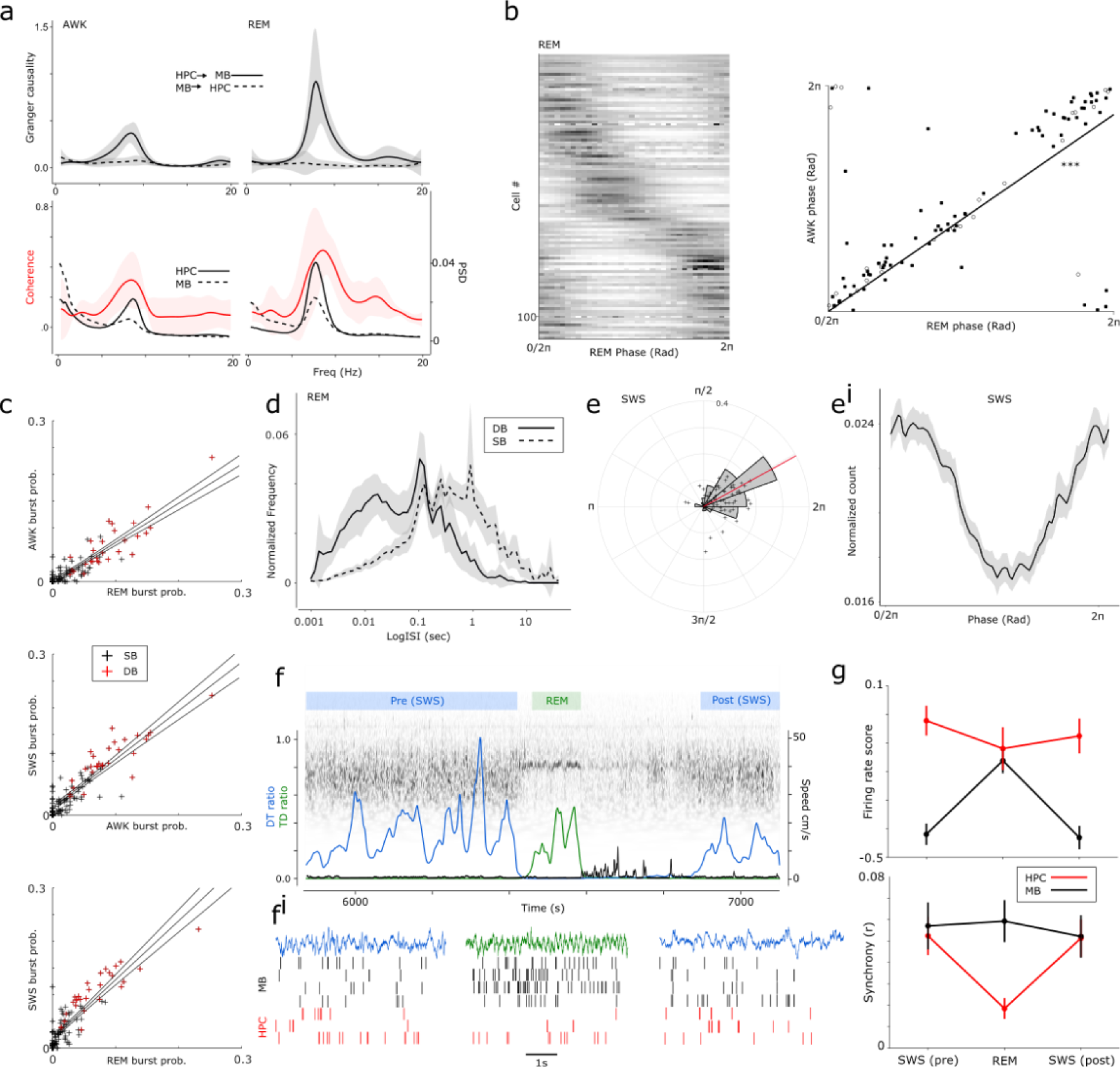
Characteristics of medial MB unit physiology across the sleep-wake cycle; **A**, In AWK and REM, both theta-dominant states, Granger causality scores (within 6-12 Hz) were significantly higher in the HPC-MB direction as compared to the reverse (upper panels). Both HPC and medial MB local field potentials (LFP) had primary peaks in the theta band with significant coherence peaks at the corresponding frequency; **B**, Units that were phase entrained during AWK, maintained their preferred phase during REM. Solid points represent sparsely bursting units and open circles, dominantly bursting units. **C**, Linear regression of bursting probability of medial MB units across sleep-wake states. Medial MB units that exhibited complex bursting in AWK, also did so during SWS and REM; **D**, Accordingly, using the classification of sparsely (SB) versus dominantly bursting (DB) bursting units defined previously for awake firing, logISI plots of DB units (see Fig. 3B) exhibited higher counts of spikes within <∼15ms relative to the logISIs of the SB units reflecting their preserved propensity to burst during REM. **E-E^i^**, Phase histogram with overlaid scatter plot of mean resultant vector length and preferred phase for each medial MB unit (**E**) alongside the mean normalized spike count phase histogram of the proportion of medial MB units (56%) that showed a phase entrainment to the down-state of 1-4Hz oscillations during SWS (slow waves; **E^i^**); **F**, Representative example of a wavelet-transformed epoch of REM sleep falling between SWS epochs, and the physiological characteristics used for REM (high theta/delta ratio; green) and SWS (high delta/theta ratio; blue), both with no movement speed; note the brief post-REM wakefulness (black line showing running speed). **F^i^**, Example 5 second epochs of wideband HPC LFP from REM (green) and neighboring SWS (blue) epochs with corresponding raster plots of synchronously recorded MB (black) and HPC PYR (red) single units; **G**, Firing rates (upper) of HPC (red) and medial MB (black) units within REM and neighboring SWS epochs, and normalized by AWK firing rates; The lower panel shows the synchrony (r) across the same sleep state transitions.

To investigate state-specific changes in activity during sleep, cell firing rates and synchrony were compared between HPC PYR and MB units during REM epochs (mean duration: 104.1 ± 2.7s) and their neighboring SWS epochs (duration: SWS pre, 600.9 ± 29.9s; SWS post, 673.6 ± 25.8s; **Fig 4F**). Comparison of cell-firing rates between HPC PYR and MB units across sleep states (15 sessions) revealed an effect of region (F_(1,14)_ = 8.02, p =0.01) and sleep state (F_(2,28)_ = 4.17, p = 0.03) on firing rate, and an interaction between region and firing rate changes across different sleep states (F_(2,28)_ = 18.27, p <0.001; **Fig 4G**). Post-hoc tests revealed lower firing rate scores for MB units during SWS epochs than REM (SWS(pre)-REM: t_(14)_ = -5.74, p <.001; SWS(post)-REM: t_(14)_ = -7.55, p <.001), and no difference between SWS epochs either side of REM (SWS(pre)-SWS(post): t_(14)_ = 0.83, n.s.). Across all sleep states, MB firing rates were lower than during AWK states (SWS(pre) : t_(16)_ = -12.10, p = < .001; REM : t_(16)_ = -3.94, p = 0.001; SWS(post): t_(16)_ = -11.35, p < .001). There was no difference between HPC firing rate scores between REM and SWS states (SWS(pre)-REM: t_(14)_ = 2.06, n.s.; SWS(post)-REM: t_(14)_ = 1.71, n.s.; SWS(pre)-SWS(post): t_(14)_ = 1.12, n.s.), however HPC firing rate during REM was lower than AWK (t_(14)_ = -3.00, p <.01), while firing rates during both SWS states were no different than AWK (SWS(pre): t_(14)_ = -1.10, p =0.29; SWS(post): t_(14)_ = -1.79, p = 0.09).

Synchrony (mean cell-pair correlations) was compared between regions and sleep states for all sessions with at least one cell pair in each region (n = 12). There was no effect of region on synchrony (F_(1,11)_ = 1.02, p = 0.33), however, there was an effect of sleep state (F_(2,22)_ = 5.25, p = 0.01), and an interaction between region and sleep-state (F_(2,22)_ = 13.00, p < 0.001). Post-hoc comparisons revealed there were no changes in synchrony in the MBs between sleep states (SWS(pre)-REM: t = -0.26, n.s.; SWS(post)-REM: t = -0.59, n.s.; SWS(pre)-SWS(post): t = 0.7, n.s.), while synchrony in HPC decreased in REM and recovered in the subsequent SWS epoch (SWS(pre)-REM: t = 4.95, p < 0.01; SWS(post)-REM: t = 4.93, p < 0.01; SWS(pre)-SWS(post): t = -0.27, n.s.). Together, these comparisons reveal a dissociation of sleep-state related modulation of activity between regions, with MB units maintaining synchrony throughout all stages of sleep, despite changes in their firing rate, and HPC PYR’s synchrony showing increases in synchrony during SWS.

Medial MB units that showed theta rhythmicity in their firing and/or theta cycle phase preference during AWK typically showed the same activity during REM. As a population, preferred firing phase, of medial MB units that exhibited significant phase preference in both states of arousal, was highly correlated between these states (circular correlation coefficient = 0.88, p <0.001). Similarly, medial MB units that showed a propensity to burst during AWK were also highly likely to burst during both SWS and REM states (REM/AWK; 0.73±0.04, t=20.30; REM/SWS 0.91±0.05, t=18.25, AWK/SWS; 1.10±0.06, t = 18.37; all p<0.001). As one would expect, given the abundance of theta rhythmicity and theta phase-entrainment in medial MB unit firing, medial MB spike - and burst-triggered averages of the HPC LFP during AWK and REM - showed medial MB spike-related epochs of high amplitude theta oscillatory activity. Meanwhile, inspection of spike-triggered averages of medial MB units in SWS epochs showed delta-dominant spike-associated activity. Finally, phase spike count histograms of SWS spikes showed that a proportion of medial MB units were significantly phase entrained during SWS (61/108 medial MB units; 56%). Furthermore, unlike in AWK and REM, at a population level, medial MB units showed a single preferred phase in SWS, firing almost exclusively during the downstate of slow waves (1-4Hz; mean resultant phase, 28.8°, Rayleigh Z =37.34, p<0.001; **Fig. 4E-E^i^**).

### Medial mammillary body unit firing is associated with hippocampal sharp wave ripples

Three recent studies have implicated the medial MBs as an important target node in the propagation of hippocampal SWR activity by the subiculum. Firstly, Kitanishi et al (2021) demonstrated that a considerable proportion of MB- but not ATN-projecting subiculum neurons fire as a function of hippocampal ripples; secondly, Kinnavane et al (2018) showed that MB-projecting vesicular glutamate transporter VGLUT2 subiculum neurons send collateral projections to the retrosplenial cortex; thirdly, Nitzan et al. (2020) demonstrated that stimulation of these VGLUT2 subicular projection neurons elicits ripple-like activity in the retrosplenial cortex. Taken together, this would suggest that at least some neurons in the medial MBs are also receiving an SWR-related input. With this in mind, we next sought to ascertain whether medial MB unit activity changes with respect to HPC SWRs during SWS. Medial MB spike-triggered averages of the hippocampal LFP during SWS revealed, in many cases, putative SWR events. The same analysis applied to the medial MB LFP showed that putative hippocampal SWRs were often associated with corresponding low frequency events in the medial MB LFP. To address whether these were, in fact, hippocampal SWRs, and medial MB response events, respectively, we next detected HPC ripple events (150-250 Hz) in the hippocampal LFP. Ripple event-triggered averages of the hippocampal LFP provided validation of our ripple-detection approach (**Fig. 5C**; red trace), with the mean average across animals showing a characteristic SWR shape. The same approach with the medial MB LFP showed that the mean response event waveform, i.e., a putative medial MB response event was remarkably consistent between animals. Temporally, the depolarization trough of the medial MB response events had a lag of approximately 40ms (lag from ripple detection: 0.042±0.001 s; **Fig. 5C**) with respect to the HPC ripple event, which would be consistent with them being temporally associated. To determine whether medial MB response events were temporally related to hippocampal SWRs, we next detected high gamma-ripple band (100-250 Hz) events in the medial MB LFP and generated peri-event time histograms of medial MB response events with respect to hippocampal SWRs. Counts of MB events within a 25 ms window around SWRs was significantly greater (p<0.01; when compared to shuffled (n=1000) ripple time distributions) in 20/32 recording sessions analysed (**Fig. 5G**). Event-triggered averages of detected MB events appeared to be highly consistent in shape with those detected by medial MB spike- and burst-triggered averages, as well as in hippocampal ripple triggered averages, of the medial MB LFP (**Fig. 5G**).

**Figure 5.**
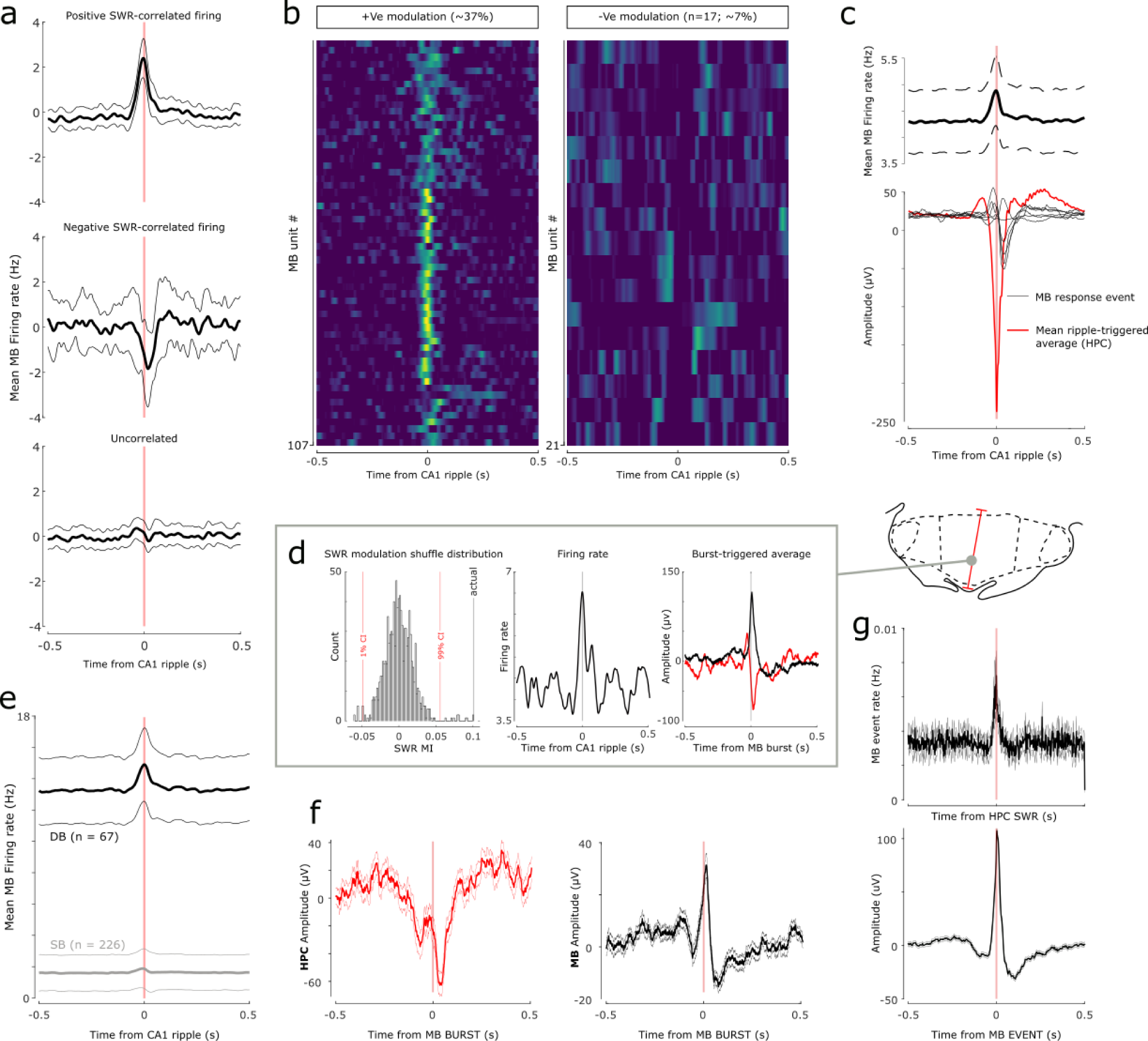
Hippocampal sharp-wave ripple (SWR)-dependent firing in the medial mammillary bodies (MBs); **A**, Mean firing rate of medial MB units in the time-window surrounding detected hippocampal SWRs. Top – middle, and lower panels include units with significant positive, negative, or uncorrelated firing, respectively; **B**, Heatmaps with each row representing the convolved mean firing rate of a medial MB unit with respect to detected HPC SWRs. Left and right panels show the (n=107 and n = 21) units with significant increases, and decreases in firing rates, respectively; **C**, HPC ripple-triggered average of the HPC (red) and the MB (black) LFPs. Individual MB black traces (n=6) represent the mean average for each animal. Upper panel shows the medial MB population-wide firing rate change with respect to HPC-detected ripples; **D**, Representative example of a medial MB unit that exhibited a significant increase in firing rate around the detected HPC ripple event times. The SWR responsiveness index (SWR RI) of the unit was higher than the 95% CI of a shuffle distribution of SWR RIs (see Methods for details). The example is also a dominantly bursting unit. Burst-triggered averages of the HPC (red) and MB (black) LFP showed a typical HPC SWR, and a medial MB response event, respectively; **E**, Mean firing rate of dominantly bursting (DB) versus sparsely bursting (SB) medial MB units, with respect to HPC ripple events. Firing rates of both DB- and SB-classified units were significantly correlated with HPC ripple events, however the mean magnitude of firing rate change was greater in the DB units. **F**, Mean average burst-triggered average of the HPC (red) and MB (black) LFP during SWS revealing large amplitude events corresponding to putative HPC SWRs and medial MB response events, respectively; **G**, Peri-event time histogram showing the mean count of MB detected events with respect to putative SWRs detected in the hippocampal LFP (150-250 Hz; Upper); Lower, Event-triggered average of high frequency (100-250 Hz) events detected in the MB LFP. The resulting waveform share the characteristics of putative MB response events observed by proximity to hippocampal SWRs (**Fig 5C**) or through burst-triggered averages of SWR responsive firing units (**Fig 5D,F**).

We next determined whether medial MB unit firing rates were correlated with ripple events. For this, we generated peri-event time histograms from which SWR modulation indices were calculated. To evaluate whether changes in MB unit were by chance, the SWR responsiveness index was compared to confidence intervals of a distribution of 999 SWR modulation indices generated from shuffle distributions based on 1000 circularly time-shifted (0-5 seconds) ripple time series (see Methods for further detail). Of 293 medial MB units recorded during SWS, 44% (125/293) of medial MB units exhibited a significant increase (n = 106; **Fig. 5A-B**, left panel), or decrease (n = 19, **Fig. 5A-B**, right panel) in firing rate. Furthermore, when cross-referenced with the bursting (DB/SB) classification outlined in Figure 3, 13% (38/293) of medial MB units exhibited conjunctive bursting and SWR-dependent changes in firing rate. Indeed, as a population, medial MB units that exhibited SWR-dependent firing had a significantly higher propensity to burst than non-SWR-dependent units (burst probability: 0.024±0.007, t = 3.504, p = 0.001 **Fig. 5D-E**).

For further evidence that medial MB DB units were exhibiting SWR-dependent firing, we generated burst center-triggered averages of all DB medial MB units during SWS. Burst-triggered averages during SWS showed the characteristics of putative hippocampal SWR events (**Fig. 5F**; see also **Fig. 5d** for representative example unit). Similarly, in the medial MB LFP, the equivalent analysis produced an event waveform that was characteristic of putative medial MB response events.

## Discussion

There have been a paucity of physiological studies focusing on the medial mammillary nuclei (medial MB), in contrast to the lateral mammillary nuclei, whose electrophysiological properties have been well-studied with respect to their role within the head direction system (Dillingham and Vann, 2019). In the present study, we classified over 500 medial MB units and found the physiological characteristics, and relative proportions of cells, to be in line with earlier reports (Sharp and Turner-Williams, 2005).

However, in addition to cells showing running speed related firing, angular head velocity, theta firing rhythmicity and theta phase preference, we report, for the first time in awake rodents, complex bursting activity and cycle skipping firing/bursting in medial MB spike trains. By recording from the medial MB across the sleep wake cycle, we found that across theta dominant states (AWK/REM), theta-related characteristics, i.e., phase preference and theta rhythmicity in spike trains of units, were largely conserved, and the bursting characteristics of cells was consistent across states, including slow-wave sleep (SWS). Finally, we report that a proportion of medial MB (37%) units exhibit hippocampal sharp wave ripple (SWR)-responsive firing (**Fig. 6**).

**Figure 6.**
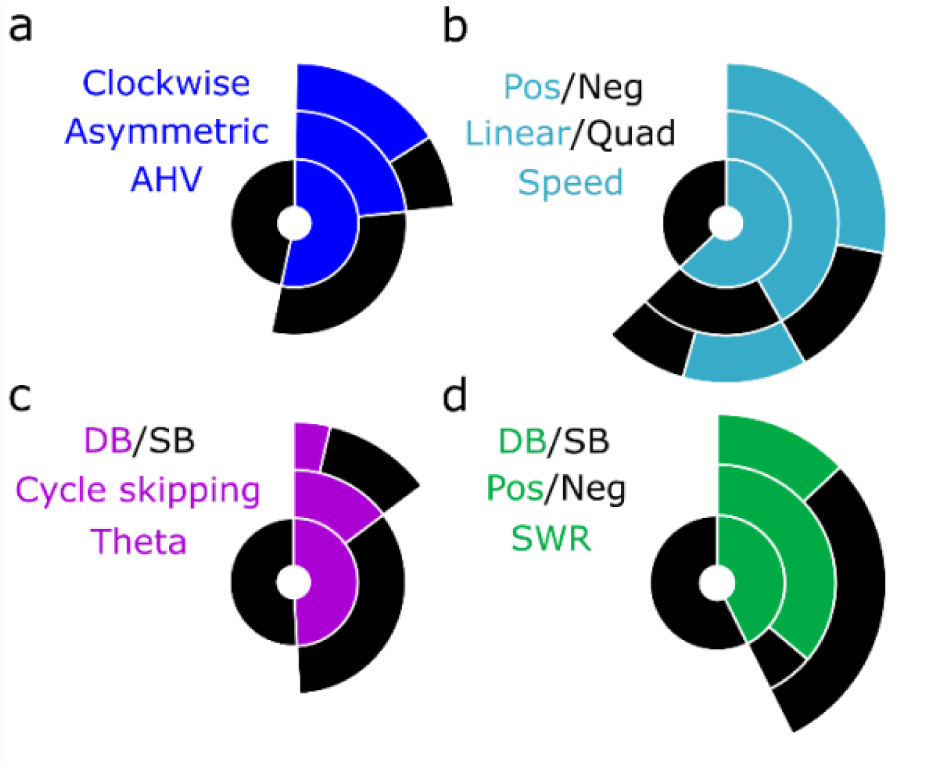
Summary charts of medial MB cell firing properties in wake and sleep. **A-D** Donut charts displaying the proportion of cells with firing rates related to the labelled property, further denoted by label color. Proportions of **A**, AHV dependent units (blue), separated by whether they are asymmetric (blue) or not (black), and the direction of movement in which asymmetric AHV units are modulated by (clockwise: blue, counterclockwise: black) **B**, speed dependent units, their best fitting polynomial (linear or quadratic fit), and the direction of the relationship between speed and firing rate **C,** theta modulated units, the presence of theta cycle skipping and the proportions of these units that are dominantly bursting (DB) or not (SB), and **D**, sharp-wave ripple (SWR) responsive units, the direction of their SWR-modulation and the proportion of SWR units exhibiting bursting.

We found similar proportions of units with running-speed dependent firing (62.8%) to Sharp and Turner-Williams (2005; 57.9%). However, we found fewer units to be positively correlated (67% vs ∼90%) and, while Sharp and Turner-Williams found speed-dependent firing in the medial MB to be linear, we found that the fit of 33.4% of units was significantly improved by inclusion of a quadratic term. A non-linear fit suggests that firing rates plateau at modest speeds in these units, similar to the saturating speed cells reported in the medial entorhinal cortex (Hinman et al., 2016). The discrepancy in findings is likely due to the speed correlations being assessed over a shorter range in the Sharp and Turner-Williams study (0-40 cm/s) making the saturation in firing rate less apparent. As a population, the running speed-responsive firing properties of medial MB units share a number of characteristics with those reported in hippocampal and parahippocampal regions (Hinman et al., 2016) and anterior thalamic nuclei (Lomi et al., 2023), i.e., in proportions of units that show either linear, or saturating speed relationships (**Fig. 2A**). How the speed signal in the medial MBs contributes to the greater speed-dependent circuitry is unclear; the MBs might passively relay the speed signal from the hippocampal formation to the anterior thalamic nuclei. However, given that the medial MBs also receive vestibular inputs via the ventral tegmental nucleus of Gudden (VTg; Irle et al., 1984), they may have a role in integrating movement-related information with theta frequency.

Approximately a quarter of medial MB units (24.7%) were found to have dominant bursting (DB) characteristics (**Fig. 6**), and the propensity for DB neurons to burst was remarkably consistent across arousal states. The majority of these bursting units also conjointly fired in relation to speed, angular velocity and/or theta. This bursting activity may be particularly important for the synchronization of activity within the medial MBs and may result in an enhanced, more reliable transmission of information to the anterior thalamic nuclei (Lisman, 1997; Csicsvari et al., 1998a; Zeldenrust et al., 2018). Spike trains can also be segregated into single spike and bursting activity, facilitating the transfer of two different information streams (Oswald et al., 2004). The presence of such bursting activity in the medial MBs raises the question of whether the medial MBs might act to segregate spatial, motor, and theta-related information, transmitting this information to the anterior thalamic nuclei through parallel information streams, or whether this bursting activity acts to ensure the stability of transfer of particularly salient information during mnemonic processing.

The medial MBs receive input from theta-dominant structures, i.e., the HPC (subiculum), the VTg, and septum (Dillingham et al., 2021). Consistent with these inputs, we found a large proportion of medial MB units (∼50%) that exhibit rhythmic firing at theta-frequencies with a broad distribution of preferred theta phases, which aligns with previous findings (Kocsis and Vertes, 1997; Sharp and Turner-Williams, 2005). However, we also observed theta cycle-skipping firing in the medial MBs. While theta cycle-skipping unit activity has been reported in the hippocampus (Harris et al., 2003), parahippocampal cortices (Deshmukh et al., 2010), prefrontal cortices (Tang et al., 2021), and nucleus reuniens (Jankowski et al., 2014), it has not previously been reported in the medial MBs. Across these different regions, theta-skipping cells have been linked to the representation of different environments and different future trajectories (Jezek et al., 2011; Kay et al., 2020; Robinson and Brandon, 2021; Tang et al., 2021). The presence of theta cycle skipping units implicates the medial MBs as a node in this prospective coding network. Given the importance of the medial MBs for spatial working memory and the acquisition of spatial tasks (Santin et al., 1999; Vann and Aggleton, 2003), it will be important to assess how theta-cycle skipping in the medial MBs contributes to these functions.

Our interest in physiological changes in the medial MBs across the sleep-wake cycle was driven primarily by recent, complementary findings on subicular-medial MB projections: 1. subicular-medial MB units show SWR-dependent firing (Kitanishi et al., 2021); 2. VGLUT2 subicular neurons send SWR information to the retrosplenial cortex (Nitzan et al., 2020); and 3, this VGLUT2 subicular-retrosplenial projection sends collaterals to the medial MBs (Kinnavane et al., 2018). Our finding, that a considerable proportion of medial MB units (∼37%) increased their firing rate around hippocampal SWR events, supports the idea that subicular projections to the medial MBs carry SWR-related information during SWS. An interesting aspect of SWR-responsive firing in the medial MBs was the combined contribution of both dominant bursting and sparsely bursting units, suggesting that the two primary efferent targets of the medial MBs, the anterior thalamus, and VTg, may be receiving multiple SWR-related information streams. Ripple-correlated bursting activity within the medial MBs, rather than isolated spikes, might also suggest that SWR-responsive activity in the medial MBs is modifying rather than simply relaying hippocampal information. As well as showing that nearly all medial MB-projecting units from the subiculum exhibit strongly SWR-responsive firing, Kitanishi et al (2021) showed that anteroventral thalamic nucleus-projecting subiculum neurons showed reduced, weak, or non-SWR-responsive firing. Anterior thalamic neurons do, however, show SWR-coupled activity (Viejo and Peyrache, 2020); the possibility exists, therefore, that propagation of hippocampal SWR-information to the anterior thalamus is dependent on the subiculum-medial MB pathway. Indeed, the medial MBs, which receive at least some of the same SWR-related input as the retrosplenial cortex, are well-placed to influence the temporal coupling of hippocampal SWRs, thalamo-cortical spindles, and delta waves in the cortex. The nesting of these events during SWS is associated with improved consolidation of hippocampal-dependent memory (Maingret et al., 2016; Kim et al., 2019), and induction of spindles through optogenetic stimulation of the thalamic reticular nucleus (with which anterior thalamic nuclei share connectivity (Lozsádi, 1995), improves hippocampal-dependent consolidation by increasing frequency of coupling of these events in the cortex (Latchoumane et al., 2017). Like the medial MB, another major hippocampal output target, the lateral septum (LS), contains a high number of SWR-responsive neurons (Wirtshafter and Wilson, 2019; Tingley and Buzsaki, 2020; Howe and Blair, 2022). In the medial MBs, high frequency (100-250 Hz) SWR response events were temporally correlated with hippocampal SWRs (**Fig. 5G**); In the LS, high frequency events, were often synchronized with SWRs but were also observed independently of HPC, and stimulation of GABAergic LS neurons elicited high frequency events locally (Tingley and Buzsaki, 2020). Unlike the LS, medial MB neurons are uniformly excitatory but in the absence of GABAergic neurons locally within the medial MBs, high frequency oscillations could feasibly be generated through recruitment of GABAergic VTg neurons via mammillothalamic axon collaterals (Hayakawa and Zyo, 1989; Umaba et al., 2021).

A second reason for assessing sleep-cycle dependent changes, was that firing rates in VTg have been reported to increase dramatically during REM (Bassant and Poindessous-Jazat, 2001) compared to during wakefulness. Given that the medial MBs have dense reciprocal connections with VTg, we were interested to see if medial MB unit firing showed similar characteristics. Our observations of firing rate changes across the sleep-cycle found the contrary. Medial MB units exhibited a small but significant reduction in firing rates during REM sleep when compared to AWK. This reduction in firing rate may reflect increased inhibition from GABAergic VTg inputs (Bassant and Poindessous-Jazat, 2001), alternatively, given that a large proportion of medial MB units exhibit motion-responsive firing (e.g. running speed and AHV), it is possible that it reflects the absence of vestibular and motion-related input during sleep. However, we also reported a greater reduction in MB unit firing rates during SWS sleep than REM sleep, thus reduction in motility alone cannot explain the extent of the changes in MB firing rates across sleep states. The modulation of MB firing rates across sleep states is in stark contrast to the relative stability of HPC PYR rates across sleep states. Sleep-state related changes in synchrony also differed between medial MBs and PYR units, with medial MB units maintaining synchrony across sleep state transitions, while PYR ensembles desynchronized during REM. Taken together with our findings that medial MB units maintain their theta phase preference during REM and exhibit phase-entrainment to the down-state of slow waves during SWS (**Fig. 4E**), it is possible that MB units are entrained to the dominant oscillatory frequency across sleep states, and changes in firing rates across sleep reflect those frequencies that define REM, and SWS. When considering that a significant proportion of MB units exhibit SWR-mediated increases in firing, the overall reduction of medial MB firing rates during SWS may provide a mechanism that reduces background activity to, in effect, amplify SWR events to the ATN. Perhaps unsurprisingly, given the size and location of the medial MBs, very few *in vivo* recording studies have focused on this region, in freely moving animals. The present study both consolidates and extends current knowledge showing how the physiological properties in the medial MBs may support their role in complex memory formation. The physiological characteristics of the medial MBs largely reflect their dominant hippocampal inputs. Strikingly, a majority of neurons exhibit conjugate firing properties, e.g., a combined running speed, angular head velocity, bursting, theta rhythmicity representation, suggesting that unlike the parallel lateral MB pathway, in which the degree of complexity in neuron representations increases from the bottom-up (Dillingham and Vann, 2019), the medial MBs are involved in higher order processing. Further functional heterogeneity is demonstrated by the varied responses of medial MB units to hippocampal SWR events, as well as widespread theta cycle-skipping activity. These combined properties point towards an active role of medial MBs in the formation of complex spatial memories (McNaughton and Vann, 2022).

## Acknowledgements

This work was funded by a Wellcome Trust Senior Research Fellowship awarded to SDV (WT212273/Z/18). The authors declare no competing financial interests. Thanks to Michal Milczarek for helpful suggestions.

## Notes

### Competing Interest Statement

The authors have declared no competing interest.

